# Astrocytic contribution to auditory hypersensitivity in a mouse model of fragile X syndrome

**DOI:** 10.1101/2025.02.14.638346

**Authors:** Lara Bergdolt, Alexandria Anding, Ola Alshaqi, Zahra Arbabi, Michael Douchey, Ember Eldridge, Katherine Hoffman, Ashley Reid, Kiran Sapkota, Ragunathan Padmashri, Daniel Monaghan, Olga Taraschenko, Anna Dunaevksy

## Abstract

Fragile X syndrome (FXS) is the most common form of inherited intellectual disability and a leading cause of autism spectrum disorder (ASD). FXS is caused by mutations in the fragile X messenger ribonucleoprotein gene 1 (*FMR1*), which result in complete or partial loss of expression of its protein product, fragile X messenger ribonucleoprotein (FMRP). Neuronal impairments in the absence of FMRP have been extensively characterized. However, much less is known about the impact that loss of FMRP has on the physiology and function of astrocytes and the implications for behavior. A common behavior exhibited by both FXS and ASD patients is hypersensitivity to sensory stimuli, but how astrocytes contribute to hypersensitivity in the context of FXS remains unknown. Using mice with astrocyte-specific reduction of *Fmr1* (*Fmr1* conditional KO (cKO)) and mice with astrocyte-specific expression of *Fmr1* (*Fmr1* cON), we demonstrated that reduction of astrocytic FMRP is sufficient but not necessary to confer susceptibility to audiogenic seizures, an indication of auditory hypersensitivity. In addition, reduction of astrocytic FMRP impacts neuronal activity, resulting in spontaneous seizures. In contrast, we assessed tactile hypersensitivity using a whisker stimulation paradigm but did not detect significant differences in *Fmr1* cKO mice. Our results reveal that astrocytes lacking FMRP contribute to auditory hypersensitivity and spontaneous seizures.

## 1. INTRODUCTION

Complete or partial loss of expression of the fragile X messenger ribonucleoprotein gene 1 (*FMR1*) causes fragile X syndrome (FXS, Pieretti et al., 1991) and is the most common single genetic aberration associated with autism spectrum disorders. Silencing of *FMR1* is caused by expansion of CGG repeats, hypermethylation of CpG islands, and histone modifications leading to chromatin condensation in the 5’ untranslated region of the *FMR1* gene (Coffee et al., 2002; Fu et al., 1991; Kremer et al., 1991; Sutcliffe et al., 1992), resulting in complete or partial loss of expression of its protein product, fragile X messenger ribonucleoprotein (FMRP). FMRP is highly expressed in the brain (Hinds et al., 1993) and regulates protein expression and function at both the mRNA and protein levels (Brown et al., 2010; Deng et al., 2013; Ferron et al., 2014; Laggerbauer et al., 2001; Li et al., 2001). Not surprisingly, loss of this regulatory protein negatively affects neurodevelopment and brain function; FXS is the most common form of inherited intellectual disability and is further characterized by social deficits, language impairments, hyperactivity, childhood seizures, and sensory hypersensitivity (Hagerman et al., 2017). Sensory hypersensitivity is thought to contribute to other impairments such as learning difficulties, anxiety, and sleep disturbances (Cascio, 2010; Sinclair et al., 2017), indicating that it is particularly important to understand the underlying mechanism(s) of this symptom.

There is increasing behavioral evidence for hypersensitivity to sensory stimuli in *Fmr1* knockout (KO) mice. A robust and often replicated indication of auditory hypersensitivity, first reported by Musumeci and colleagues (Musumeci et al., 2000), is susceptibility to audiogenic seizures. Brief exposure to a sustained loud sound often causes tonic-clonic seizures in *Fmr1* KO mice. Tactile hypersensitivity is also observable in *Fmr1* KO mice in multiple assays including a gap-cross assay (Arnett et al., 2014; Juczewski et al., 2016), which requires that mice use their whiskers to gather information about their surroundings, and a whisker stimulation assay, during which unilateral whisker stimulation induces more aversive and defensive behavior in *Fmr1* KO mice (He et al., 2017; Kourdougli et al., 2023). In a novel assay designed to evaluate the response of mice to social touch, *Fmr1* KO mice also exhibited more avoidance and aversive behavior towards social touch than WT controls (Chari et al., 2023). *Fmr1* KO mice also exhibit impaired discrimination in learning paradigms that depend on sensory (visual, auditory, and olfactory) processing in the presence of sensory distractors (Kuruppath et al., 2023; Rahmatullah et al., 2023). Several indications of neuronal hyperexcitability have been associated with auditory and tactile stimulation throughout development and into adulthood (Arnett et al., 2014; He et al., 2017; Jonak et al., 2024; Rotschafer & Razak, 2013; Zhang et al., 2014). In addition, neurons in the somatosensory cortex of *Fmr1* KO mice exhibit increased UP state duration (Gibson et al., 2008; Hays et al., 2011) and elevated firing rate (Goncalves et al., 2013), further supporting the hyperexcitability of neurons in the absence of FMRP.

Astrocytes, a type of glial cell in the brain, may contribute to these phenotypes, as they are known to regulate neuronal signaling at the cellular and circuit levels. For example, astrocytes are involved in the regulation of UP states in neurons of the somatosensory cortex (Poskanzer & Yuste, 2011), state switching in cortical circuits (Poskanzer & Yuste, 2016), and neuronal synchronization in the hippocampus (Sasaki et al., 2014). Yet, astrocytes have not been studied as extensively as neurons in the context of FXS. Astrocytes express FMRP in multiple brain regions throughout development (Gholizadeh et al., 2015; Higashimori et al., 2016), suggesting that silencing of *Fmr1* in astrocytes can contribute to the phenotypes of FXS. Indeed, deletion of FMRP in mouse astrocytes has been shown to alter dendritic morphology (Jacobs & Doering, 2010; Jacobs et al., 2010; Yang et al., 2012), cause differential expression of proteins that regulate synaptic formation (Cheng et al., 2016; Krasovska & Doering, 2018; Wallingford et al., 2017), impact density of dendritic spines and colocalization of synaptic puncta (Cheng et al., 2016; Higashimori et al., 2016; Hodges et al., 2016; Jacobs et al., 2010; Krasovska & Doering, 2018; Wallingford et al., 2017), impair glutamate transport and signaling (Higashimori et al., 2013; Higashimori et al., 2016), increase duration of neuronal UP states (Jin et al., 2021), enhance neuronal excitability (Bataveljic et al., 2024), and contribute to behavioral impairments including learning, memory, and social interaction (Hodges et al., 2016; Jin et al., 2021). Notably, stem cell-derived astrocytes from individuals with FXS, when co-cultured with human stem cell-derived neurons, also contribute to abnormal neuronal firing (Sharma et al., 2023), indicating the translational relevance of studying the effects of *Fmr1* deletion in mouse astrocytes.

However, the contribution of astrocytes to sensory hypersensitivity in FXS remains unknown. Here we employed two assays for sensory hypersensitivity that have been characterized in *Fmr1* KO mice to determine the astrocytic contribution to these phenotypes. Specifically, we tested susceptibility to audiogenic seizures (AGS), an indication of auditory hypersensitivity, and response to repeated whisker stimulation, an indication of tactile hypersensitivity. We further investigated the effect that loss of astrocytic FMRP has on neuronal activity *in vivo*, both at baseline using electroencephalogram (EEG) recordings and following acoustic stimulation by analyzing c-Fos density. Overall, our study identifies astrocytes as contributors to auditory hypersensitivity in the absence of FMRP.

## 2. METHODS

### 2.1 Mice

Mice were cared for in accordance with NIH guidelines for laboratory animal welfare. All experiments were approved by the Institutional Animal Care and Use Committee at the University of Nebraska Medical Center. Mice were housed in ventilated cages with *ad libitum* access to food and water. Female breeders were given ENVIGO-Teklad 2019 rodent diet to promote lactation and litter survival and post-weaning experimental mice were given ENVIGO-Teklad 8656 or later 7012 when 8656 was no longer available. Mice were maintained on a 12-hour light-dark cycle. All experiments were performed during the light phase except for EEG recordings which were continuous throughout the light and dark phases.

Most experiments were performed using male mice because the *Fmr1* gene is on the X chromosome and FXS is more common in males than females. We included a few female mice in the whisker stimulation experiments and saw no obvious differences between males and females of the same genotype. We used the following mouse lines: Aldh1l1-Cre/ERT2 (Srinivasan et al., 2016, Jackson #029655), GCaMP6f^flox^ (Madisen et al., 2015, Jackson 024105), tdTomato^flox^ (Madisen et al., 2010, Jackson #007908), *Fmr1* KO (Consortium, 1994, Jackson #003025), *Fmr1*^flox^ and *Fmr1*^loxP-neo^ (from Dr. David Nelson; Guo et al., 2011; Mientjes et al., 2006). All mice were maintained on a C57BL/6 background, although Aldh1l1-Cre/ERT2 mice still contain traces of FVB alleles (Srinivasan et al., 2016). *Fmr1* KO and wild type littermate (WTLM) experimental mice were generated by crossing *Fmr1*^+/-^ females to males that were *Fmr1*^-/y^, Aldh1l1-Cre^+^/GCaMP^f/f^, or *Fmr1*^+/y^ (to generate WTLM females). Mice that expressed Aldh1l1-Cre and GCaMP6f received tamoxifen following the same injection protocol as the *Fmr1* cKO mice (see below) and were grouped with *Fmr1* KO and WTLM for the audiogenic seizure assay. To drive astrocyte-specific conditional deletion (*Fmr1* cKO) or expression (*Fmr1* cON) of *Fmr1*, we used the tamoxifen-inducible Cre line Aldh1l1-Cre/ERT2. Aldh1l1-Cre^+^/tdTomato^f/wt^ or Aldh1l1-Cre^+^/GCaMP6f^f/wt^ mice were used to determine recombination efficiency in different cell types following early postnatal tamoxifen injections. *Fmr1* cKO experimental mice and littermate controls were generated by breeding Aldh1l1-Cre^+^/Fmr1^f/wt^ females with males that were Aldh1l1-Cre^+^, Aldh1l1-Cre^+^/GCaMP6f^f/f^, or Aldh1l1-Cre^+^/*Fmr1*^f/y^ (to generate cKO females). These crosses yielded *Fmr1* cKO (Cre^+^/*Fmr1*^f/y^, Cre^+^/*Fmr1*^f/f^) and WTLM (Cre^+^/*Fmr1*^wt/y^, Cre^+^/*Fmr1*^wt/wt^, Cre^-^/*Fmr1*^wt/y^, Cre^-^/*Fmr1*^wt/wt^, Cre^-^/*Fmr1*^f/y^, or Cre^-^/*Fmr1*^f/f^). *Fmr1* cON experimental mice and littermate controls were generated by breeding *Fmr1*^loxP-neo/wt^ or *Fmr1*^loxP-neo/loxP-neo^ females with either Aldh1l1-Cre^+^ or Aldh1l1-Cre^+^/GCaMP6f^f/f^ males. These crosses yielded *Fmr1* cON (Cre^+^/*Fmr1*^loxP-neo/y^), WTLM (Cre^+^/*Fmr1*^wt/y^ or Cre^-^/*Fmr1*^wt/y^), and knockout littermates (KOLM; Cre^-^/*Fmr1*^loxP-neo/y^).

Tamoxifen (Sigma T5648) was resuspended in corn oil (Sigma C8267) and was administered to all mice, regardless of Cre expression, according to the following paradigm: 50 μL of 1 mg/mL tamoxifen injected in the milk sac on postnatal days (P) 1-3 and 20 mg/kg (using 2 mg/mL tamoxifen), intraperitoneal (i.p.) injection on P5 and P7.

### 2.2 Tissue fixation and immunohistochemistry

Recombination efficiency in astrocytes, neurons, and oligodendrocytes in Aldh1l1-Cre mice following tamoxifen administration was determined on P17-21 as follows. Brains of mice expressing Aldh1l1-Cre and GCaMP6f or tdTomato were fixed via cardiac perfusion with 4% paraformaldehyde, extracted, and postfixed overnight in 4% paraformaldehyde before storage in 1X PBS at 4°C. Coronal sections 100 μm thick were cut using a vibratome (Leica VT1000S). For immunohistochemistry, sections were washed three times (5 min. each) with 1X PBS, incubated with 0.3% Triton (Fisher) in PBS for 3 min., then blocked for one hour with 5% normal goat serum (NGS, Vector) in 0.3% Triton, all at room temperature with gentle shaking. Next, sections were incubated with primary antibodies in 5% NGS plus 0.3% Triton overnight at 4°C with gentle shaking. Primary antibodies used were anti-S100β (1:250; Abcam ab868 or 1:1000 Synaptic Systems 287004), anti-NeuN (1:500, Millipore MAB377), anti-Olig2 (1:500, Millipore ab9610), and anti-GFP (1:500; Invitrogen A10262) to visualize GCaMP6f. The following day, sections were washed 3 times (10 min. each) with PBS and incubated with secondary antibodies in 1% NGS plus 0.3% Triton. Secondary antibodies used were Alexa Fluor 594 anti-mouse (Invitrogen A11005), Alexa Fluor 594 anti-guinea pig (Invitrogen A11076), Alexa Fluor 647 anti-rabbit (Invitrogen A21245), Alexa Fluor 647 anti-mouse (Invitrogen A21235), Alexa Fluor 488 anti-rabbit (Invitrogen A11008), and Alexa Fluor 488 anti-chicken (Invitrogen A11039) at half the concentration that was used for the primary antibody. Sections were washed 3 times (10 min. each) and mounted on microscope slides with Fluoro-Gel mounting medium (Electron Microscopy Sciences). Images were acquired with a Nikon A1R upright confocal microscope, 20X objective (0.75 NA) with a resolution of 512 x 512 (pixel = 0.62 µm). Image stacks with a step size of 1 µm were collected from the primary motor or somatosensory cortex (1200 μm or 2500 μm from the midline), with 2-4 images acquired per section. Maximum intensity projected images of 5 optical sections were created in ImageJ, which was also used for analysis of recombination efficiency to determine the proportion of S100β+, NeuN+, or Olig2+ cells that are also tdTomato+ or GFP+ (GCaMP6f+).

For analysis of FMRP levels in astrocytes following tamoxifen administration, brains of *Fmr1* cKO, *Fmr1* KO, and WTLM mice that expressed GCaMP6f were fixed on P17-21 as described above. Brains were subsequently sectioned and processed for immunohistochemistry as described above, with an extra blocking step to prevent the anti-mouse IgG secondary antibody from binding to endogenous mouse proteins. Specifically, following blocking with 5% NGS, sections were washed with PBS 3 times (5 min. each), sections were blocked with 1X ReadyProbes^TM^ Mouse-on-Mouse Blocking Solution (Invitrogen R37621) for one hour, then washed with PBS 3 times (5 min. each), followed by incubation with primary antibodies in 5% NGS overnight. The protocol for the second day is the same as above. Primary antibodies used were anti-FMRP (1:100; BioLegend 834701), anti-S100β (1:1000 Synaptic Systems 287004), and anti-GFP (1:500; Invitrogen A10262) and secondary antibodies were Alexa Fluor 594 anti-mouse (Invitrogen A11005), Alexa Fluor 647 anti-guinea pig (Invitrogen A21450) and Alexa Fluor 488 anti-chicken (Invitrogen A11039). Sections were mounted on microscope slides with Fluoromount-G with DAPI (Invitrogen 00-4959-52). Images were acquired as above, except with a 60X objective (1.40 NA) at a resolution of 1024 x 1024 (pixel = 0.21 µm) and a step size of 0.5 µm. Analysis in Fiji was performed on summed intensity projections of 5 optical frames, centered around the middle of astrocyte soma. S100β labeling was used to identify the middle of the soma of astrocytes in layer I of the cortex and subsequently to perform manual outlining of astrocyte soma and any processes that could be seen in the projected image. The mean gray value of FMRP in a small region away from somata was considered to be background and was used to perform background subtraction on all FMRP images before measuring the mean gray value in astrocytes.

For analysis of c-Fos+ cells in *Fmr1* cKO and WTLM following auditory stimulation, mice were exposed to a loud alarm as for the audiogenic seizure test (described below). Only mice that did not exhibit seizure behavior were used for c-Fos analysis. Sixty minutes following siren exposure, brains were fixed, sectioned, and immunostained as described above, with the following modifications. The density of c-Fos+ cells was examined in the midbrain (inferior colliculus, IC), auditory thalamus (medial geniculate body, MGB), and the primary auditory cortex (A1) to determine if there are regional differences in c-Fos expression between the WTLM and cKO mice. Sagittal brain sections were used for IC and coronal brain sections were used for MGB and A1. Two-three sections for each region of interest were used for c-Fos immunohistochemistry for each mouse (n=7-10 mice). Sections were chosen based on the Allen mouse brain atlas for consistency in slice locations across mice. Consistency in selecting the auditory cortex sections was based on the location relative to the hippocampus (Martin del Campo et al., 2012). For immunohistochemistry, c-Fos (Cell Signaling 2250) and NeuN (Millipore MAB377) primary antibodies were used at 1:500 and secondary antibodies (Alexa Fluor 488 anti-rabbit, Invitrogen A11008 and Alexa Fluor 647 anti-mouse, Invitrogen A21235, respectively) were used at 1:1000.

Stained sections were imaged using a confocal microscope as described for recombination efficiency above, except with a 10X objective and a step size of 2 µm. All images of the same staining were acquired using the same settings. Image analysis was performed on maximum intensity projected images (5 optical sections) using ImageJ software. The dimensions of the region of interest used for cell counting were based on the auditory structure analyzed (IC: manual tracing of the entire IC; MGB: 300 x 600 µm; A1: 400 µm wide outline spanning cortical layers II/III-VI). A rolling ball background subtraction was done for all images. The c-Fos+ cell counts were based on intensity autothresholding of pixels (methods used: triangle for IC, Otsu for MGB, moments for A1) and size (Analyze Particles) in ImageJ. The watershed function was applied to each image to separate overlapping cells.

### 2.3 Audiogenic seizures

Susceptibility to audiogenic seizures (AGS) was determined as previously described (Westmark et al., 2011). Male mice were tested on P18-23, the age of peak susceptibility to AGS in *Fmr1* KO mice on a C57BL/6 background (Yan et al., 2004). Mice were brought to the testing room at least 30 minutes before testing for acclimation. Testing was performed between 12:00 and 19:00. Five minutes before testing, the mouse was placed in a plexiglass chamber (20 x 7.5 x 20 cm) with a personal alarm (LOUD KEY^TM^) present but not on for additional acclimation and video recording of baseline behavior. Testing consisted of exposure to the alarm (118-122 dB) for five minutes. Sound intensity was confirmed with a sound meter each day before starting experiments. Each session was video recorded for subsequent analysis. The total number of mice exhibiting seizure behavior was scored. Seizure behavior always began with wild running and often progressed to tonic-clonic seizures and sometimes death. The average seizure score for each group was determined by assigning a seizure score to each mouse (Tao et al., 2023) as follows: 0 = no seizure behavior, 1 = one bout of wild running, 2 = two bouts of wild running, 3 = one tonic-clonic seizure, 4 = two tonic-clonic seizures, 5 = death. Latency and duration of seizure behavior were used as additional measures of seizure properties.

### 2.4 Cycloheximide treatment

Cycloheximide (Tocris) was prepared fresh daily at a concentration of 10 mg/mL in 0.9% NaCl. A dose of 120 mg/kg or equivalent volume of vehicle was administered i.p. one hour before AGS testing.

### 2.5 Implantation of EEG electrodes

Male mice at 5-6 weeks old were anesthetized with isoflurane and implanted with a 2-EEG/1EMG head mount and two cortical screw EEG electrodes (Pinnacle Technology, Lawrence, KS) positioned to record signals from the parietal cortex overlying the posterior hippocampus (EEG 1) and ipsilateral frontal cortex (EEG 2). The reference screw electrode was positioned over the contralateral parietal cortex. The depth of anesthesia during surgery was assessed by tail pinch. Following the surgery, animals were housed individually and monitored daily; postoperative analgesia was achieved with carprofen (5 mg/kg i.p. daily for 2 days). Following 7-9 days of recovery, mice were placed in the EEG recording chambers and connected to the acquisition system.

### 2.6 Assessment of audiogenic seizure susceptibility in EEG electrode implanted mice

To determine if exposure to the siren caused seizures that did not manifest in behavior and to measure neural oscillations, we performed electroencephalogram (EEG) recordings on 6-7 week old *Fmr1* cKO and WTLM. Following the recording of baseline EEG for 20-24h, mice were transferred to a sound-attenuating custom-made chamber equipped with a video camera with infrared capability and reconnected to the EEG acquisition system. Following habituation for one hour, a strobe siren (126 dB; Fortress) was activated for 5 min. Time of siren activation was recorded to enable later identification of EEG segments corresponding to sound delivery.

### 2.7 EEG acquisition and analysis

The EEG system (Pinnacle 8206, Pinnacle Technology) was comprised of a preamplifier unit connected by a tether to a conditioning/acquisition system. Signals were sampled at 400 Hz (preamplifier gain at 100X, total gain 5,000X, high pass EEG channel filter: 0.5 Hz, low pass EEG filter: 50 Hz), digitized using a 14-bit analog to digital converter and routed to a PC.

EEG analysis was carried out using Sirenia Seizure Pro 1.7.6 (Pinnacle Technology Inc.) and verified visually without knowledge of the genotype. The EEG signal was categorized as normal or seizures (greater than 5 sec of sustained, rhythmic high amplitude activity in both EEG leads) as previously reported (Taraschenko et al., 2019; Taraschenko et al., 2021). The duration of each seizure, total seizure count, and time to the first seizure from the start of EEG acquisition was recorded for each mouse. The corresponding video was reviewed during and 10 sec after electrographic seizure candidate events to characterize seizure behavior.

Two 5-min long segments of baseline EEG recordings during artifact-free, awake activity were combined for analysis of neural oscillations. EEG data were divided into 1 sec segments and Fast Fourier Transforms (FFT) algorithm were used on each segment. The average power was calculated for the delta (0-4 Hz), theta (4-8 Hz), alpha (8-13 Hz), beta (13-30 Hz), and gamma (30-50 Hz) bands for each electrode and compared between *Fmr1* cKO and WTLM mice.

### 2.8 Telemetry surgery

Surgical implantation of a subcutaneous biotelemetry transmitter (Data Sciences International (DSI) PhysioTEL HD-X02/S02), which is outfitted with EEG and EMG leads, was performed at P60-P90. To reduce inflammation, carprofen (5 mg/kg, i.p.) was administered 15 minutes prior to the procedure. Mice were anesthetized with 2% isoflurane, shaved from the head to upper back, then head-mounted onto a stereotaxic frame. During surgery, a heating pad was used to maintain body temperature and eye lubricant was applied to the eyes to prevent drying. Following administration of bupivacaine (1 mg/kg, s.c.), a 3 cm midline incision was made from the posterior boundary of the eye to mid-scapulae level. Two EEG leads were secured through bilateral burr holes drilled into the skull (1.5 mm lateral and 3.0 mm posterior to Bregma, 1.0 mm lateral and 1.0 posterior to Lambda). Two EMG leads were secured in nuchal muscle. The transmitting device was inserted into a subcutaneous pocket along the left flank between the forelimb and hind limb. After surgery, carprofen was administered for seven days, enrofloxacin (5 mg/kg, i.p.) was administered for 10 days, and bacitracin was applied topically to the incision site for seven days. Mice were housed individually during recovery and for the remainder of the experiment.

### 2.9 Telemetry recording and seizure analysis

Each cage was placed on a telemetry receiver in a dedicated room in the Animal Behavior Core at the University of Nebraska Medical Center. Mice were allowed to acclimate for 4 hours, and a continuous 24-hour recording was executed with bandpass filter of 0.1-1000 Hz and sampling rate of 500 Hz using the DSI Ponemah software. In addition to the EEG and EMG recordings, the physical activity of the mouse was recorded both by video and the telemetry device. Lights turn on at 06:00 and turn off at 18:00. Animals had access to food and water *ad libitum*. EEG analysis was carried out using DSI Neuroscore software (v. 3.4) and verified visually, using the same criteria as described above, except there was only one EEG lead for these recordings.

### 2.10 Whisker stimulation and assessment of avoidance behavior

Avoidance behavior during tactile stimulation was assessed by analyzing the response of head-restrained mice to repeated whisker stimulation. On P28-31, stainless-steel headplates for restraining the mouse were implanted as follows. To reduce inflammation, carprofen (5 mg/kg) was administered 15 minutes prior to the procedure and once daily for three days following headplate placement. The mouse was anesthetized with isoflurane and its head was secured using a stereotaxic setup on top of a warm water circulating pad to maintain body temperature. Puralube vet ointment (Dechra) was applied to the eyes to prevent corneal drying. Breathing and toe pinch reflex were monitored during the procedure and the amount of isoflurane administered was adjusted accordingly. Fur on the top of the head was shaved, and the area was disinfected with povidone-iodine and ethanol wipes before the skin was removed. Xylocaine (Fresenius Kabi) was applied to the skull and a scalpel was used to scrape off connective tissue. The skull was cleaned with acetone and quickly dried. A modified headplate (Neurotar, Model 1 with the posterior two blades cut off to prevent ears from getting stuck in clamp) was glued to the skull above lambda with Krazy glue. A mixture of DuraLay powder (Reliance) and Loctite 401 was used to secure the headplate and cover all areas of exposed skull. Following the procedure, 200 μL of saline was administered subcutaneously to aid in recovery.

Mouse handling and habituation to the Mobile HomeCage (Neurotar) were performed as previously described (Kislin et al., 2014; Padmashri et al., 2021). Briefly, mouse handling was performed the day after headplate implantation. First, the experimenter, wearing terrycloth autoclave gloves, allowed each mouse to freely crawl around the experimenter’s hands for 3 sessions lasting approximately 3 minutes each. The interval between sessions varied but each session was at least 10 minutes apart. Next, the mouse was securely wrapped in a small cloth three times with varying intervals between sessions. Habituation to the Mobile HomeCage began later that day or the following day and consisted of one 20-minute session each day for 10 days. Lights were on for the first session and off for all subsequent habituation sessions as well as during testing. A custom-ordered carbon fiber cage with a 1 cm wall was used to prevent interference with the filaments used for whisker stimulation.

The whisker stimulation experiment was performed as previously described with modifications (He et al., 2017). Testing was performed between 13:00 and 17:00. We used the Mobile HomeCage with locomotion tracking which detects the mouse’s position in the cage and its trajectory. An infrared camera was used to record each trial to identify when filament movement started and ended as well as for confirmation of the locomotion tracking data. The whisker stimulation setup (Figure 5A) consisted of a comb of six filaments approximately 1.5 mm apart descending from bent glass capillaries, which were attached to a piezoelectric actuator (PiezoDrive PD200). The stimulation protocol consisted of 20 stimulations lasting 1 s each, with a 3 s interstimulus interval. Before testing, the mouse was given three minutes to acclimate to the Mobile HomeCage. Next, a 3-minute baseline was recorded. Immediately following the baseline, a sham recording was performed during which the stimulation filaments were moving but were positioned just out of reach of the whiskers on the left side. Performing the sham stimulation first allowed us to isolate the response of each mouse to the sound that accompanies movement of the filaments before the mouse had a chance to associate this sound with tactile stimulation. For the stimulation, the filaments were intercalated with the whiskers approximately 3 mm from the skin on the left side of the mouse’s nose.

For locomotion tracking in the Mobile HomeCage, sensors in the air dispenser track the position of two magnets in the cage and that data is extrapolated to yield the position, velocity, and trajectory of the mouse. Data was acquired at a rate of 77 or 84 frames per second, depending on the software version. To determine the total movement of the mouse, we calculated the amount of time the mouse was moving at or above 8 mm/s. We further divided total movement in four ways depending on timing and direction: movement during or between stimuli and movement towards or away from stimulus. Timestamps from the start of stimulation and from each frame of the motion tracking data were used to identify segments during stimulation or interstimulus intervals. Avoidance (movement “away” from stimulus) and non-avoidance (movement “towards” stimulus) during stimulation were determined by the change in alpha (Δα = α_n_ – α_n-1_), where α is defined as the angle between the mouse’s longitudinal axis and the Y-axis of the cage. Positive Δα indicates the cage is moving counterclockwise, as if the mouse is trying to move to the right. Conversely, negative Δα indicates the cage is moving clockwise, as if the mouse is trying to move to the left. Since the filaments for stimulation are in contact with the whiskers on the left of the mouse’s snout, we considered positive Δα to indicate avoidance behavior and negative Δα to indicate non-avoidance. For both total movement and avoidance movement, we calculated a habituation score by taking the ratio of average movement during the last five stimulations to average movement during the first five stimulations, such that a habituation score of less than one indicates less movement during later trials compared to earlier trials, which we interpret as habituation to the repeated stimuli. Mice that did not move during the first five trials were excluded from habituation analysis as we need a non-zero value for initial movement to calculate the habituation ratio.

### 2.11 Statistics

Whisker stimulation sham and stimulation comparisons were analyzed by repeated measures ANOVA using SAS 9.4. Statistical analyses of all other data were performed using GraphPad Prism (version 10). Normality of all continuous data was tested using the Shapiro-Wilk test. Outliers were identified using the ROUT method and were removed from all datasets. Unpaired t-tests were used for normally distributed continuous data (with Welch’s correction when group variances were significantly different) and Mann-Whitney tests were used for non-normally distributed continuous data and categorical data. Kruskal-Wallis test was used for non-normally distributed data with three or more groups. Chi-square tests were used to compare frequency distributions.

## 3 RESULTS

### 3.1 Generation of experimental Fmr1 cKO and cON mice

To isolate the effect of astrocytic FMRP on sensory hypersensitivity, we utilized mice with conditional deletion (*Fmr1*^flox^) or expression (*Fmr1*^loxP-neo^) of *Fmr1* (Mientjes et al., 2006). These mice were bred with Aldh1l1-CreERT2 mice to generate experimental mice, which we refer to as *Fmr1* conditional KO (cKO) for astrocyte-specific deletion of *Fmr1*, and *Fmr1* conditional on (cON) for astrocyte-specific expression of *Fmr1*. We chose the Aldh1l1-CreERT2 line because Aldh1l1 is expressed by nearly all astrocytes (Cahoy et al., 2008). In addition, the Aldh1l1-CreERT2 line is more specific for astrocytes over other cell types, compared to other astrocytic Cre lines such as GFAP-Cre or GLAST-Cre (Srinivasan et al., 2016). Since FMRP is highly active during development, we wanted to activate Cre recombinase to induce *Fmr1* deletion or expression early in development. To achieve this, we administered tamoxifen on postnatal days (P) 1-3, 5, and 7 (Figure 1A). This is the first time this tamoxifen injection strategy has been used with Aldh1l1-Cre, so we first characterized the efficiency with which Cre-induced recombination occurred in different cell types in the brain by using *Fmr1* cKO mice with Cre-dependent expression of the genetically-encoded calcium indicator GCaMP6f or reporter mice with Cre-dependent expression of the fluorescent protein tdTomato. We determined that administration of tamoxifen five times within the first postnatal week yields expression of the fluorescent reporter in approximately 85% of S100β-positive astrocytes in the cortex at P17-21. Recombination occurs almost exclusively in astrocytes, with expression in only 0.1% of NeuN+ neurons and 0.4% of Olig2+ oligodendrocytes (*F*(2, 11) = 3811, *p* <.0001; Figure 1B-D). To test if expression of exogenous fluorescent reporters is an acceptable proxy for deletion of endogenous FMRP, we quantified fluorescent intensity of FMRP in S100β-positive astrocytes of P17-21 mice and normalized to fluorescence in KO astrocytes (Figure 1E). As expected, we found significantly greater fluorescent intensity in astrocytes from WTLM, compared to *Fmr1* KO mice (*H* = 34.07, *p* <.0001; Figure 1F). In addition, relative FMRP expression in GCaMP-positive astrocytes from *Fmr1* cKO mice was significantly lower compared to WT astrocytes (*H* = 34.07, *p* =.014). If a complete knockout of FMRP was achieved, we would expect relative FMRP expression in GCaMP-positive astrocytes in *Fmr1* cKO mice to be similar to that in astrocytes from *Fmr1* KO, but in fact, it is significantly higher (*H* = 34.07, *p* =.010), suggesting that we are more accurately modeling astrocytic *knockdown* of FMRP, rather than complete astrocyte-specific knockout.

**Figure 1:**
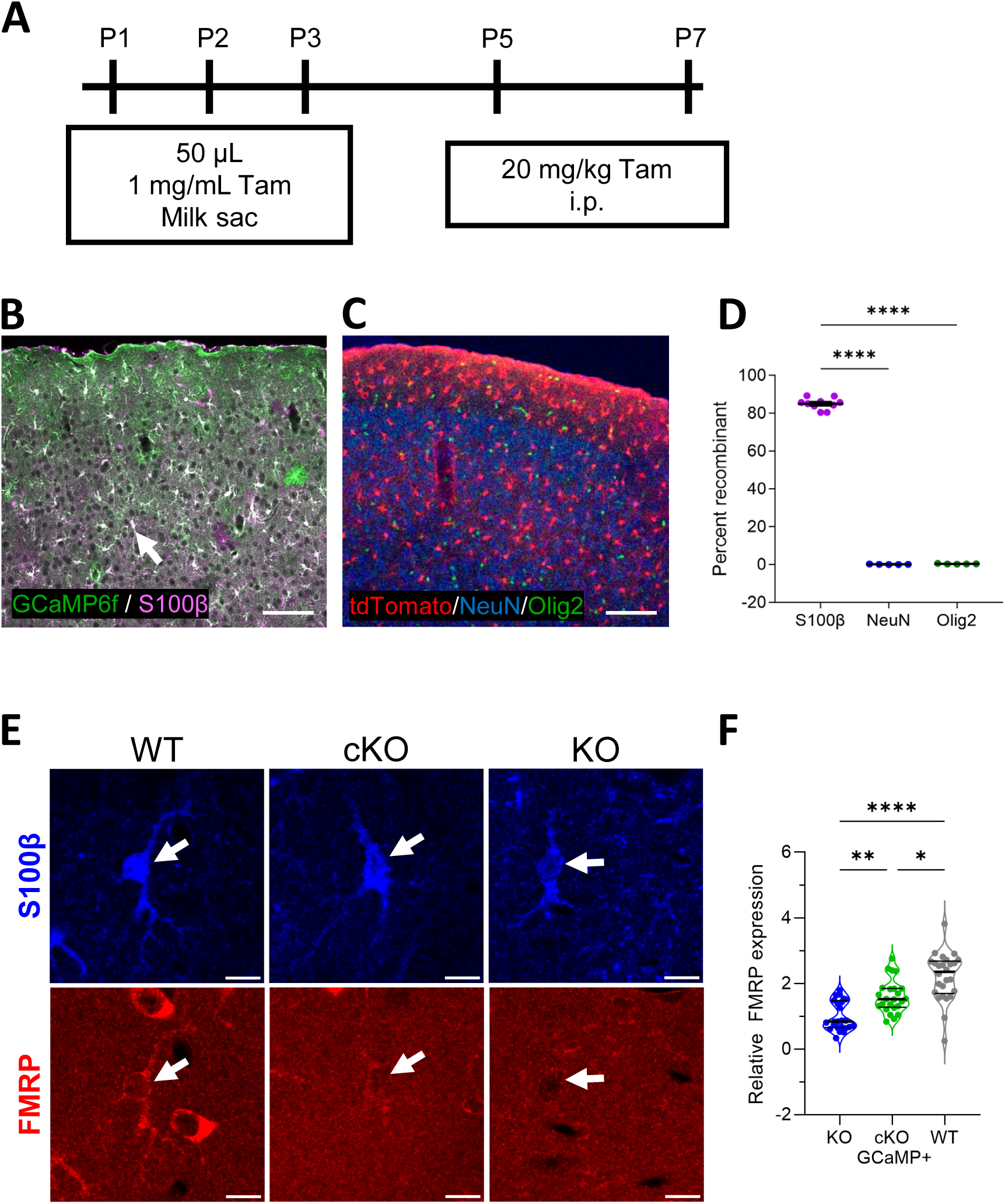
Characterization of recombination in Aldh1l1-cre mice. **A)** Timeline for tamoxifen injections. **B)** Representative image of the cortex used to determine recombination efficiency in astrocytes following tamoxifen injections. Cells in which recombination occurred express GCaMP6f (green). Astrocytes are labeled with S100β (magenta). S100β+ cells expressing GCaMP6f appear white (arrow). **C)** Representative image of the cortex used to determine recombination efficiency in neurons (NeuN+; blue) and oligodendrocytes (Olig2+; green) following tamoxifen injections. Cells in which recombination occurred express tdTomato (red). Scale bar is 100 μm. **D)** Recombination efficiency in the cortex of Aldh1l1-Cre mice (n = 5-10 mice, average of 8 images per mouse). **E)** Representative images of FMRP intensity in astrocytes from WT, *Fmr1* cKO, and *Fmr1* KO mice. Scale bar is 10 µm. **F)** Relative FMRP expression in cortical astrocytes, normalized to *Fmr1* KO (n = 26-28 images; data points are averages of 3-10 astrocytes per image, with 6-8 images acquired per mouse, 4 mice of each genotype). Data were analyzed using one-way ANOVA with Tukey post hoc tests (**D**) and Kruskal-Wallis with Dunn post hoc tests (**F**). *p <.05, **p <.01, ****p <.0001

### 3.2 Reduced astrocytic FMRP is sufficient to increase audiogenic seizure severity

Sensory hypersensitivity, including auditory hypersensitivity, is a common characteristic of FXS (Baranek et al., 2008; Butler et al., 1991; Rotschafer & Razak, 2014; Sinclair et al., 2017). *Fmr1* KO mice also exhibit auditory hypersensitivity (Lovelace et al., 2018; Lovelace et al., 2016; Wen et al., 2018). One of the ways this is expressed is as a robust phenotype of increased susceptibility to audiogenic seizures (AGS, Chen & Toth, 2001; Musumeci et al., 2000; Yan et al., 2004). Exposure to a loud siren induces seizure behavior characterized by wild-running, which often progresses to tonic-clonic seizures and sometimes death. Because astrocytes are implicated in modulation of neuronal activity and may contribute to seizure onset or progression (Vezzani et al., 2022), we determined if altered astrocytic function due to reduction of FMRP contributes to audiogenic seizure susceptibility.

Similar to previous studies (Musumeci et al., 2000; Yan et al., 2004), exposure to a 120 dB siren induced seizure behavior in a greater proportion of three-week old *Fmr1* KO mice than in WTLM (X^2^(1) = 4.95, *p* =.026; Supplementary figure 1A). We scored the severity of seizure behavior in each mouse on a scale from 0-5, with zero being no seizure behavior and five indicating death following seizure. The average seizure score is significantly higher for *Fmr1* KO than WTLM (*U* = 369.5, *p* =.014). To further characterize properties of AGS, we analyzed latency to onset of seizure behavior and total duration of seizure behavior. We use the term seizure behavior to include wild running and tonic-clonic seizure, so the duration of seizure behavior is the time from onset of wild running to the end of tonic-clonic seizure or death, when applicable. We found that seizure behavior commences after 25.4 ± 6.5 (WT) and 33.1 ± 9.7 (KO) seconds of siren exposure and continues for 34.6 ± 13.8 (WT) and 28.5 ± 4.7 (KO) seconds, but there is not a significant difference between genotypes (Latency: *U* = 24, *p* =.56; Duration: *t*(16) = 0.55, *p* =.59; Supplementary figure 1B).

To determine if reduced astrocytic FMRP contributes to AGS susceptibility, we tested three-week old *Fmr1* cKO mice, the age of peak susceptibility to AGS in *Fmr1* KO mice. As outlined in the methods, there are multiple combinations of *Fmr1* and Aldh1l1-Cre genotypes that result in mice having WT levels of *Fmr1* expression. Since there is evidence for increased susceptibility to seizures with expression of Cre under two other promoters (Emx1-Cre, expressed predominantly in excitatory neurons, and Dlx5/6-Cre, expressed primarily in inhibitory neurons) following administration of the proconvulsant compound pentylenetetrzol (Kim et al., 2013), we tested whether Aldh1l1-Cre alone impacts AGS susceptibility. We did not observe a difference in the proportion of Cre-positive and Cre-negative mice that exhibited seizure behavior or in the overall seizure score (Proportion: X^2^(1) = 0.65, *p* =.42; Score: *U*= 99, *p* =.62; Supplementary figure 2A). Based on these data, we concluded that Aldh1l1-Cre does not enhance susceptibility to AGS and therefore we combined results from mice with the following genotypes for the WTLM group: Cre^+^/*Fmr1*^wt/y^, Cre^-^/*Fmr1*^wt/y^, and Cre^-^/*Fmr1*^f/y^. Some mice also expressed GCaMP6f to facilitate analysis of Cre-mediated recombination. We did not find a significant effect of GCaMP expression on AGS susceptibility (Proportion: X^2^(1) = 0.65, *p* =.42; Score: *U* = 177, *p* =.62; Supplementary figure 2B), so all mice expressing GCaMP were grouped with their respective *Fmr1* genotype.

We observed seizure behavior in 45% of *Fmr1* cKO and in only 20% of WTLM. Although there is not a statistically significant difference in the proportion of *Fmr1* cKO mice that exhibited seizure behavior (X^2^(1) = 2.85 *p* =.091), the majority of *Fmr1* cKO mice with seizure behavior had tonic-clonic seizures while the seizure behavior of WTLM consisted primarily of wild running only, resulting in a significantly higher seizure score in *Fmr1* cKO mice (*U* = 138, *p* =.044; Figure 2A). In addition, the total duration of seizure behavior tended to be longer in *Fmr1* cKO mice than in WTLM (*t*(10) = 2.11, *p* =.061; Figure 2B), but there was no difference between genotypes in the latency to onset of seizure behavior (*U* = 12.5, *p* =.59). These results suggest that reduced astrocytic FMRP is sufficient to elevate AGS severity.

**Figure 2:**
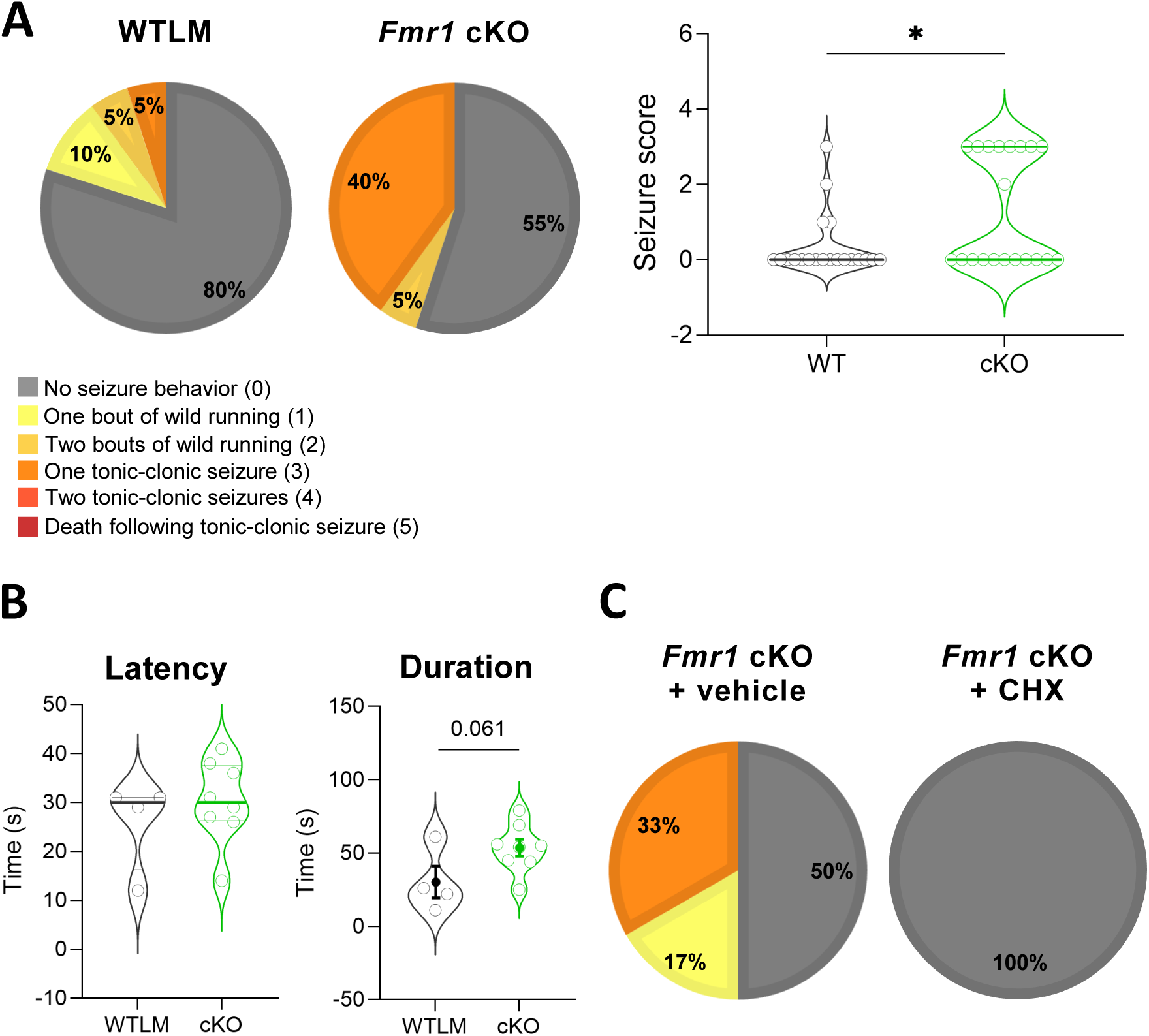
Increased susceptibility to audiogenic seizures in *Fmr1* cKO mice. **A)** Proportion of *Fmr1* cKO and WTLM mice exhibiting seizure behavior and corresponding average seizure score (n = 20 WTLM and 20 cKO). **B)** Latency to onset of seizure behavior and total duration of all seizure behavior in mice that exhibited seizure behavior (n = 4 WT and 8 cKO). **C)** Proportion of *Fmr1* cKO exhibiting seizure behavior 1 hour after treatment with the protein synthesis inhibitor cycloheximide (CHX, 120 mg/kg; n = 6 vehicle and 16 CHX). Chi-square tests were used to analyze proportion of mice exhibiting seizures (**A** and **C**); Mann-Whitney tests were used to analyze seizure score (**A**) and latency (**B**); unpaired t-test was used to analyze seizure duration (**B**). In **A** and **B** (latency), horizontal lines represent median and upper and lower quartiles. In **B** (duration), filled circles represent mean and bars represent ± SEM. *p <.05.

Following exposure to a loud auditory stimulus, *Fmr1* KO mice have a greater number of activated neurons, indicated by expression of c-Fos, in the inferior colliculus and regions of the medial geniculate body (Nguyen et al., 2020). To determine if astrocytes contribute to this phenotype, we used immunohistochemistry to examine the number of cells expressing c-Fos in a subset of *Fmr1* cKO and WTLM mice that did not exhibit AGS behavior during siren stimulation. We used mice that did not exhibit seizure behavior to avoid potential confounds from motor activity. One hour after auditory stimulation, brains were fixed via cardiac perfusion and processed for immunohistochemistry. We focused on three regions involved in binaural processing in the auditory processing pathway: the inferior colliculus, the thalamus, and the auditory cortex (Supplementary figure 3). In both the inferior colliculus and the auditory cortex, the density of c-Fos+ cells was comparable between genotypes (inferior colliculus: *t*(11) = 0.53, *p* =.61; auditory cortex: *t*(11) = 0.76, *p* =.46), although there was considerable inter-mouse variability in both genotypes. We also observed variability in the density of c-Fos+ cells in the thalamus, but there is a trend towards increased density in the thalamus of *Fmr1* cKO mice (*t*(13) = 1.95, *p* =.073), suggesting that reduced astrocytic FMRP may cause increased neuronal activity in parts of the auditory pathway following auditory stimulation.

### 3.3 Increased protein synthesis contributes to increased propensity for AGS

FMRP regulates protein expression and excess protein synthesis has been reported in the absence of FMRP (Dolen et al., 2007; Osterweil et al., 2010; Qin et al., 2005). Therefore, we wanted to determine if enhanced protein synthesis resulting from reduction of *Fmr1* in astrocytes contributes to AGS susceptibility. Treatment with cycloheximide (CHX), a protein synthesis inhibitor, significantly reduced AGS susceptibility in Fmr1 KO mice (Stoppel et al., 2021). We treated *Fmr1* cKO with CHX or vehicle and found that inhibition of protein synthesis completely prevents AGS in *Fmr1* cKO mice (X^2^(1) = 9.26 *p* =.002; Figure 2C), suggesting that the pathogenic proteins are rapidly turned over proteins that are not replenished within the one-hour timeframe.

### 3.4 Loss of astrocytic FMRP is not necessary for audiogenic seizure susceptibility

To determine if loss of astrocytic FMRP is necessary for AGS susceptibility, we tested three-week old *Fmr1* cON and KOLM mice. We found that 39% of *Fmr1* cON mice exhibited seizure behavior, compared to 48% of KOLM, indicating similar seizure susceptibility in the two groups (X^2^(1) = 0.34 *p* =.56; Figure 3A). In addition, the number of mice exhibiting each degree of seizure behavior was similar in each genotype, resulting in similar seizure scores in both genotypes (*U* = 282, *p* =.79). Seizure onset time and total duration also were not different between genotypes, indicating that restoration of astrocytic FMRP cannot prevent seizures or reduce seizure severity (Latency: *t*(19) = 0.31, *p* =.76; Duration: *U* = 26, *p* =.077; Figure 3B). Therefore, we conclude that although reduction of astrocytic FMRP is sufficient to cause AGS, it is not necessary. Rather, loss of FMRP in astrocytes may decrease the threshold for seizure onset, while not being the main driver of seizure behavior in *Fmr1* KO mice.

**Figure 3:**
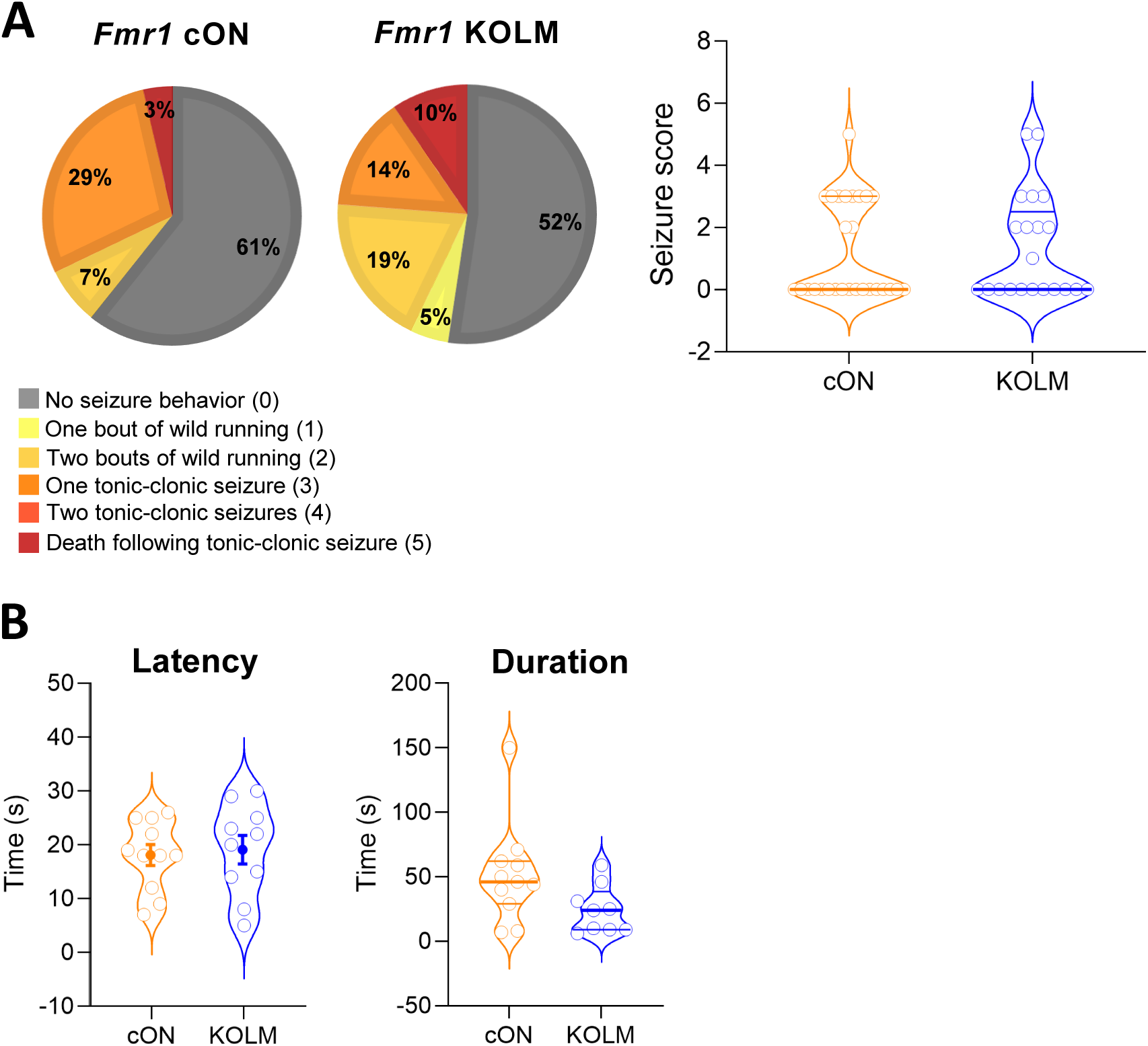
Restoration of astrocytic FMRP does not decrease AGS susceptibility. **A)** Proportion of *Fmr1* cON mice and WTLM exhibiting seizure behavior and corresponding average seizure score (n=28 cON and 21 KOLM). **B)** Latency to onset of seizure behavior and total duration of all seizure behavior (n = 11 cON and 10 KOLM). Chi-square test was used to analyze proportion of mice exhibiting seizures (**A**); Mann-Whitney tests were used to analyze seizure score (**A**) and duration (**B**) and an unpaired t-test was used to analyze seizure latency (**B**). In **A** and **B** (duration), horizontal lines represent median and upper and lower quartiles. In **B** (latency), filled points represent mean and bars represent ± SEM.

### 3.5 Spontaneous electrographic seizures in *Fmr1* cKO mice

Not all seizures are observable by behavior; many seizures can only be detected by electroencephalogram (EEG) recordings. To determine if *Fmr1* cKO mice exhibit electrographic seizures at baseline or during exposure to a 120 dB siren, we implanted EEG electrodes, positioned to record from the parietal and frontal cortices, on 5-6 week old *Fmr1* cKO and WTLM mice. We chose this age as it is the minimum age when mice consistently survive implantation of the EEG electrodes. After 7-9 days of recovery, baseline activity was recorded for 24 hours. Interestingly, half of *Fmr1* cKO mice had at least one electrographic seizure during this period, but no seizures were observed in WTLM (*U* = 18, *p* =.029; Figure 4). The total duration of all seizures combined ranged from 0.5 to 7.5 minutes for *Fmr1* cKO, with an average seizure duration of 2.0 minutes. These seizures were electrographic only; no behavioral seizures were observed at any point. Following the baseline recording, mice were transferred to a dark sound-attenuating box, allowed to acclimate for one hour, then exposed to the siren for five minutes. None of the mice exhibited seizure behavior or electrographic seizures during the 5-minute exposure to a siren. We also assessed the presence of spontaneous seizures in a cohort of freely moving adult (2-5 months) *Fmr1* KO, cKO, and WTLM mice (n = 15 KO, 14 cKO, and 13 WT) using EEG recordings from telemetry devices. We did not observe seizures in any mice at these older ages (data not shown). Thus, we conclude that while reduction of astrocytic FMRP does not increase the propensity for audiogenic seizures in adolescent *Fmr1* cKO mice, nor spontaneous seizures in adult mice, it does cause spontaneous electrographic seizures in adolescent mice.

**Figure 4:**
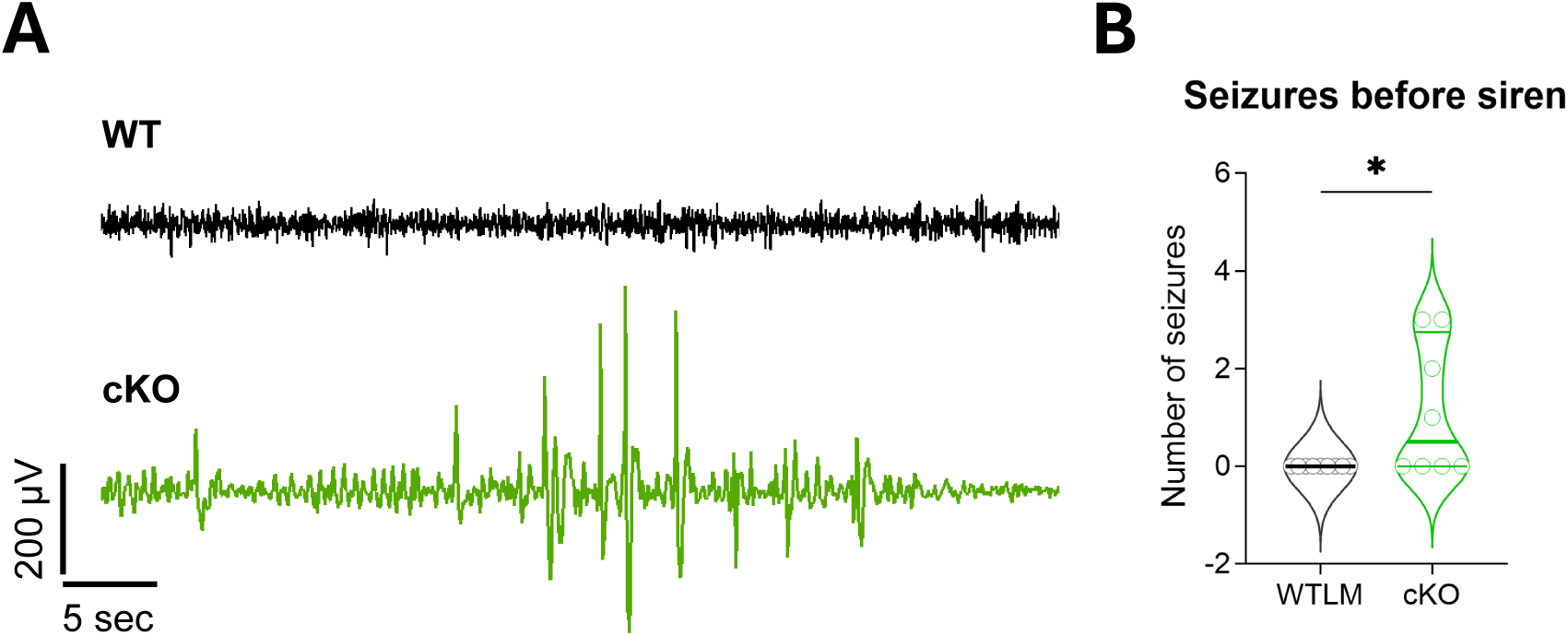
***Fmr1* cKO mice exhibit an increased number of electrographic seizures. A)** Representative EEG traces from a WT and *Fmr1* cKO mouse, illustrating an electrographic seizure in the *Fmr1* cKO mouse. **B)** Number of electrographic seizures (n=9 WT and 8 cKO, Mann-Whitney test, horizontal lines represent median and upper and lower quartiles). *p <.05

**Figure 5:**
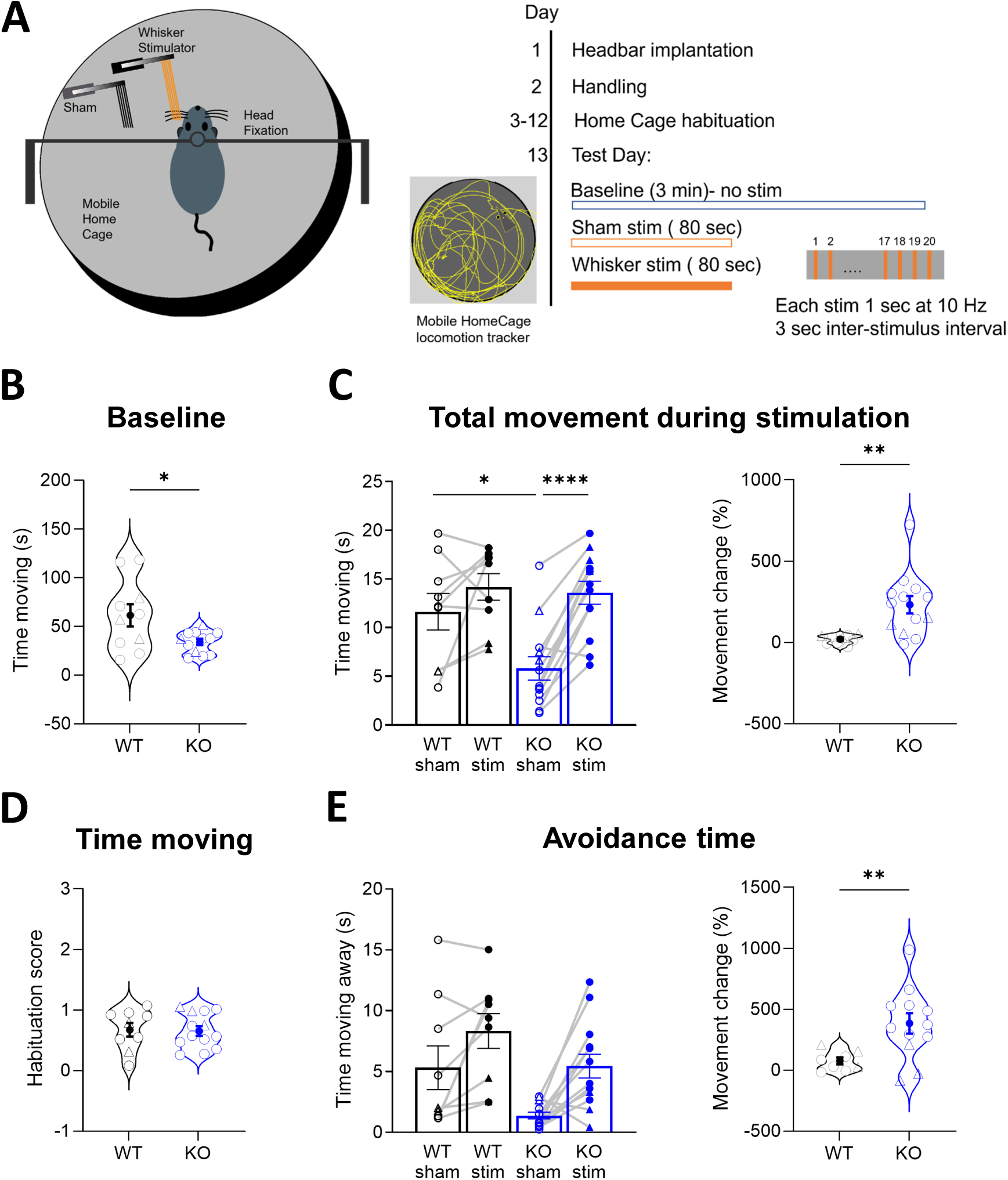
***Fmr1* KO mice exhibit tactile hypersensitivity. A)** Diagram of setup for whisker stimulation experiment and timeline of the experiment. A sample trace of movement detected by the Mobile HomeCage locomotion tracking software is included with the timeline. **B)** Total time spent moving during the baseline recording (n = 10 WT, 12 KO). **C)** Total time spent moving (left) and percent change in movement from sham trial to stimulation trial (right) during all 20 sham and whisker stimulations for *Fmr1* KO and WTLM (n=9 WT, 13 KO). **D)** Habituation score (average of last five stimulations/average of first five stimulations) for time moving (n=9 WT, 13 KO). **E)** Duration (left) and percent change from sham trial to stimulation trial (right) of avoidance movement (away from the stimulating filaments). For **B-E**, Circles represent male mice and triangles represent female mice. For **C** and **E**, gray lines connect data points from the same mouse and bars represent mean ± SEM. For **B**, **C** (right), **D**, and **E** (right), filled circles represent mean and bars represent ± SEM. Data with four groups in **C** and **E** (left) were analyzed using repeated measures ANOVA; unpaired t-tests were used for two group comparisons in in **B**, **C** (right), **D**, and **E** (right). \**p* <.05, \*\**p* <.01, \*\*\*\**p* <.0001

Given that *Fmr1* KO mice exhibit enhanced power of low and high frequency oscillations (Lovelace et al., 2018) and astrocytes modulate neural oscillations (Bellot-Saez et al., 2018; Poskanzer & Yuste, 2016), we used our EEG recordings from 6-7 week old mice to determine if reduction of astrocytic FMRP impacts neural oscillations. We sampled two segments from before presentation of the siren during which the mouse was awake. Power values were normalized by expressing mouse averages as a ratio of WT for each frequency range to allow for comparison of relative changes across all frequency ranges. However, we did not observe differences between *Fmr1* cKO and WT in the average power of any of the frequency bands in these segments (Parietal cortex: *F*(4, 63) = 2.17, *p* =.082; Frontal cortex: *F*(4, 68) =.35, *p* =.84; Supplementary figure 4).

### 3.6 *Fmr1* cKO mice do not exhibit tactile hypersensitivity

In addition to auditory hypersensitivity, *Fmr1* KO mice also exhibit tactile hypersensitivity. *Fmr1* KO mice exhibited impaired sensory processing when whiskers were stimulated (Arnett et al., 2014; Juczewski et al., 2016). In addition, compared to WTLM, adolescent *Fmr1* KO mice exhibited greater avoidance behavior by steering away from the stimulated side when whiskers were stimulated at regular intervals (He et al., 2017). We performed a similar experiment to determine if reduction of astrocytic FMRP contributes to this phenotype. We used the Mobile HomeCage (MHC) with locomotion tracking capabilities, in which the mouse is head-restrained and positioned on a flat air-lifted cage (Figure 5A). The mouse can easily move the air-lifted cage, thus giving it the illusion of exploring the cage. As hitching the mouse to the headclamp can be a stressful experience and the MHC is a novel environment for the mouse, each mouse was habituated to the setup for 20 minutes every day for 10 days, starting when mice were P29-32.

We chose to habituate for 10 days because corticosterone levels are reported to plateau after 10 days of head-fixing and habituation (Juczewski et al., 2020). On the day of the experiment, baseline movement was recorded for three minutes, followed by a sham stimulation trial, during which the stimulation filaments were positioned out of reach of the mouse’s whiskers. The sham trial was immediately followed by a stimulation trial, during which filaments were intercalated with the mouse’s whiskers, approximately 3 mm from the left side of the snout. The stimulation apparatus consisted of a comb of six filaments attached to a piezoelectric actuator that moved the filaments in the anterior-posterior direction at a frequency of 10 Hz for one second with 3 second interstimulus intervals.

Since we used a different apparatus than what has been previously described to measure avoidance behavior, we first tested a cohort of *Fmr1* KO and WTLM to confirm that *Fmr1* KO mice exhibit greater avoidance behavior with our setup and that we can detect it with the locomotion tracking software. We primarily tested male mice but also included a few female mice. To assess whether *Fmr1* KO mice exhibit differences in overall movement in the MHC, we analyzed the amount of time mice spent moving during the baseline recording. We found that *Fmr1* KO mice move significantly less than WTLM (*t*(10.5) = 2.31, *p* =.042; Figure 5B). We next compared the responses of *Fmr1* KO and WTLM mice to sham and whisker stimulation. We found a significant interaction between genotype and trial type in the total time moving (*F*(1, 20) = 7.63, *p* =.012; Figure 5C), as well as an increase in movement during stimulation relative to sham in *Fmr1* KO mice (*F*(1, 20) = 41.25, *p* <.0001) with no difference in amount of movement during sham and stimulation conditions in WTLM (*F*(1, 20) = 3.07, *p* =.095). We observed lower movement during sham trials in *Fmr1* KO mice relative to WTLM (*t*(20) =-2.75, *p* =.012), consistent with pre-stimulation baseline movement. To allow us to consider the magnitude of the change in movement from sham to stimulation trials and account for varying amounts of movement during sham trials, we calculated the percent change in movement and, consistent with our ANOVA results, found that *Fmr1* KO mice exhibit a greater increase in movement during stimulation than WTLM (*t*(12.7) = 3.91, *p* =.002). Greater movement during whisker stimulation suggests that *Fmr1* KO mice exhibit greater aversion to whisker stimulation.

Another measure that could indicate hypersensitivity is the degree of habituation to repeated stimuli. To assess habituation in our mice, we calculated a habituation score by taking the ratio of average movement during the last five stimulations to average movement during the first five stimulations. For both genotypes, the average habituation score was less than or close to one, indicating that most mice of both genotypes moved less (or a similar amount) during the end of the trial relative to the beginning of the trial (*t*(20) = 0.17, *p* =.87; Figure 5D). Thus, all mice display a similar habituation response to repeated stimuli, regardless of genotype.

A more specific measure of avoidance of tactile stimulus would incorporate the direction of the mouse’s intended movement, with attempted movement away from the stimulator considered to be avoidance. To determine if *Fmr1* KO mice also exhibit greater avoidance movement, we compared the total time of avoidance movement during sham and stimulation trials. Although there is nothing to avoid during the sham trial (the stimulation filaments are not touching the mouse), we use the term avoidance to indicate direction of movement to maintain consistency between the trial types, with avoidance indicating the cage is moving counterclockwise. We found significant main effects of both trial type (*F*(1, 20) = 21.03, *p* <.001) and genotype (*F*(1, 20) = 6.69, *p* =.018; Figure 5E) for avoidance time. Additionally, the magnitude of increase in avoidance movement from sham to stimulation was significantly higher in *Fmr1* KO, further supporting the conclusion that *Fmr1* KO mice exhibit greater aversion to whisker stimulation. In summary, we conclude that *Fmr1* KO mice demonstrate an exaggerated response to tactile stimuli in our setup, confirming that we can detect hypersensitivity to tactile stimuli.

We next used *Fmr1* cKO mice to test the specific effect of reduced astrocytic FMRP on the tactile hypersensitivity we observed in *Fmr1* KO mice. In contrast to *Fmr1* KO mice, *Fmr1* cKO mice did not exhibit a difference in baseline movement (*U* = 79, *p* =.41; Figure 6A). Additionally, we did not find a significant interaction of trial type and genotype when looking at total movement during stimulation, nor a significant main effect of genotype (*F*(1, 25) = 0.03, *p* =.86), but there was a main effect of trial type (*F*(1, 25) = 34.7, *p* <.0001; Figure 6B). Consistently, *Fmr1* cKO and WTLM mice show a similar percent increase in movement from sham to stimulation trials (*U* = 66, *p* =.54). As with the *Fmr1* KO cohort, habituation during stimulation trials does not differ between genotypes (time moving: *t*(25) = 0.25, *p* =.80; Figure 6C).

**Figure 6:**
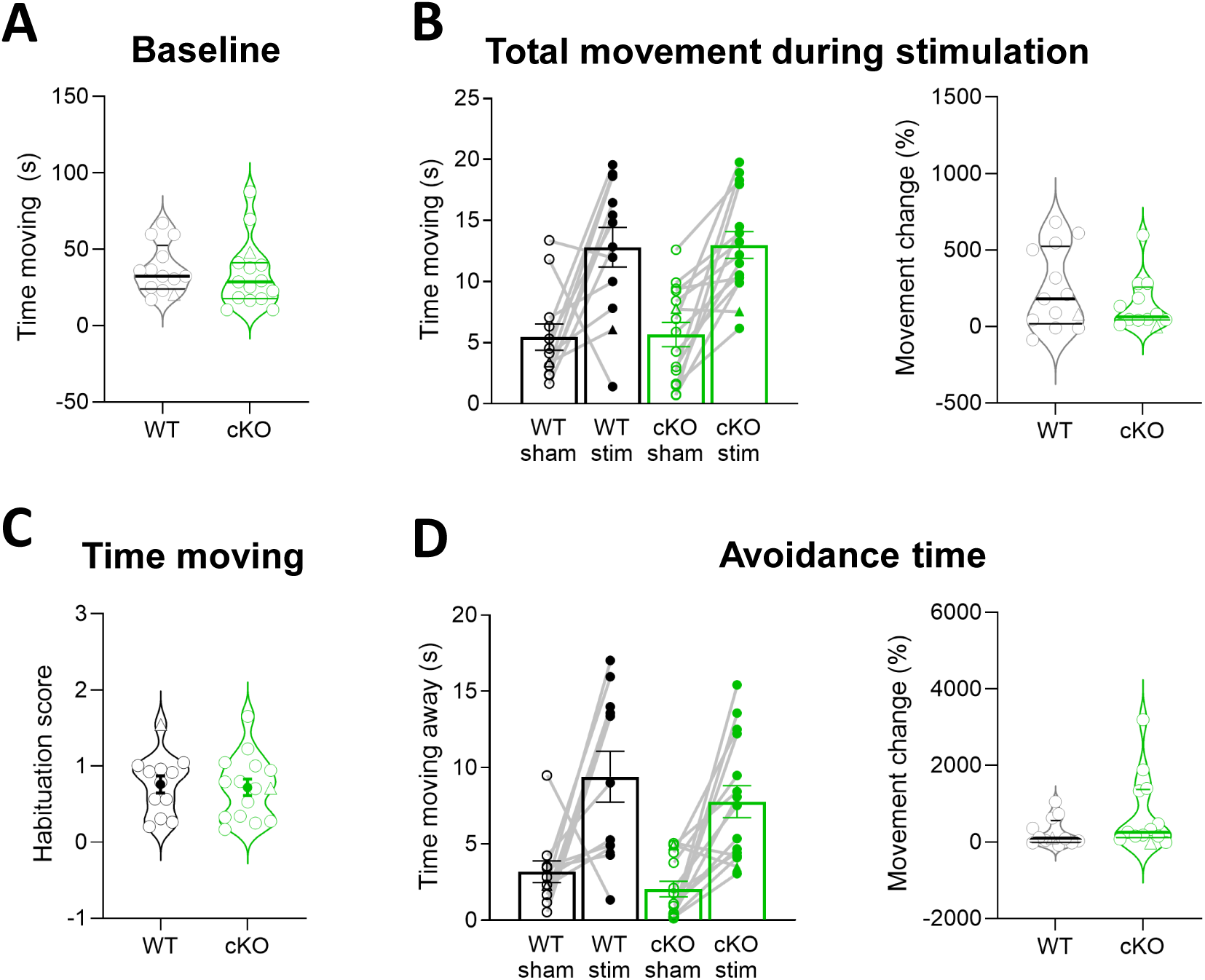
***Fmr1* cKO mice do not exhibit tactile hypersensitivity. A)** Total time spent moving during the baseline recording (n = 13 WT, 15 cKO). **B)** Total time spent moving (left; n = 12 WT, 15 cKO) and percent change in movement from sham trial to stimulation trial (right; n = 13 WT, 12 cKO) during all 20 sham and whisker stimulations for *Fmr1* cKO and WTLM. **C)** Habituation score (average of last five stimulations/average of first five stimulations) for time moving during stimulations (n = 12 WT, 15 cKO). **D)** Duration (left) and percent change from sham trial to stimulation trial (right) of avoidance movement (away from the stimulating filaments). In all graphs, circles represent male mice and triangles represent female mice. For **B** and **D**, gray lines connect data points from the same mouse and bars represent mean ± SEM. For **A**, **B**, and **D** (right), horizontal lines represent median and upper and lower quartiles. For **C**, filled circles represent mean and bars represent ± SEM. Data with four groups in **B** and **D** (left) were analyzed using repeated measures ANOVA; Mann-Whitney tests were used for two group comparisons in **A**, **B** (right), and **D** (right); unpaired t-tests were used for **C**.

For time spent in avoidance movement, we did not find a significant interaction of trial type and genotype by repeated measures two-way ANOVA, nor a significant main effect of genotype (*F*(1, 24) = 3.98, *p* =.057). However, there was a main effect of trial type (*F*(1, 24) = 27.2, *p* <.0001; Figure 6D), with both *Fmr1* cKO and WTLM moving more during stimulation. In addition, there was no difference between genotypes in the relative change in movement from sham to stimulation trials (*U* = 50, *p* =.14). In summary, no significant differences were detected between *Fmr1* cKO and WTLM mice in this whisker stimulation assay, suggesting that partial loss of astrocytic FMRP expression does not contribute to the form of tactile hypersensitivity measured here.

## 4 DISCUSSION

Using mice with specific reduction and restoration of FMRP expression in astrocytes, here we report that loss of astrocytic FMRP contributes to auditory hypersensitivity in mice. We also found spontaneous seizures detected by EEG in adolescent *Fmr1* cKO mice. However, we did not detect tactile hypersensitivity in *Fmr1* cKO mice.

The audiogenic seizure assay taps into two characteristics of FXS patients: seizure susceptibility and auditory hypersensitivity. Approximately 10-15% of children with FXS have seizures (Hoffmann & Berry-Kravis, 2016). The age-dependent decline in audiogenic seizure susceptibility in mice mirrors the timeframe of seizure occurrence in FXS patients, with seizure onset and resolution usually occurring during childhood (Berry-Kravis et al., 2021; Musumeci et al., 2000). However, a drawback to using the AGS assay as a seizure model is that seizures in FXS patients have not been reported to be triggered by loud stimuli. As such, AGS testing is better suited as a probe for auditory hypersensitivity. Our results from *Fmr1* cKO and cON mice indicate that loss of astrocytic FMRP is sufficient, but not necessary, to increase susceptibility to AGS. Sufficiency is demonstrated with a higher prevalence of seizures and an increased seizure score in *Fmr1 c*KO compared to WTLM. Restoration of FMRP expression only in astrocytes did not impact AGS susceptibility, indicating that loss of astrocytic FMRP is not necessary for increased AGS susceptibility.

*Fmr1* deletion in VGLUT2-expressing cells has been shown to be both necessary and sufficient for the AGS phenotype (Gonzalez et al., 2019) while another study observed sufficiency of *Fmr1* deletion in Purkinje cells (Gibson et al., 2023). Given the extensive crosstalk between neurons and astrocytes, it is conceivable that loss of FMRP in both cell types contributes to the hyperexcitability that leads to AGS susceptibility, especially considering electrophysiology recordings in slices from astrocyte-specific *Fmr1* cKO mice revealed evidence for neuronal hyperexcitability in the form of elevated NMDAR-mediated EPSC amplitude and longer spontaneous UP states (Jin et al., 2021). It is also plausible that there was some astrocytic contribution when *Fmr1* was deleted in VGLUT2-expressing cells, as there is now evidence for expression of VGLUT2 in a subset of astrocytes (de Ceglia et al., 2023). The fact that restoration of *Fmr1* expression in two distinct neuronal populations reduces AGS susceptibility while restoration in astrocytes has no impact on AGS behavior suggests that reduction of astrocytic FMRP may lower the threshold for seizure onset.

We observed AGS behavior in 41% of *Fmr1* KO and 48% of KOLM from cON litters. This is lower than what has been previously reported for *Fmr1* KO mice on a C57BL/6 background, which ranges from approximately 60-75% for 3-week old mice (Gross et al., 2019; Osterweil et al., 2013; Sawicka et al., 2016; Westmark et al., 2011; Yan et al., 2004). One possible explanation for the lower prevalence is the level of ambient noise mice are exposed to in the caging system. Our mice were housed in microisolator cages with mechanical ventilation, making the housing environment louder than it would be with static isolator cages and perhaps increasing the threshold of sound tolerance. However, the fact that we observed higher AGS scores in *Fmr1* KO mice than in WTLM, indicates that, even being housed with higher ambient noise levels, *Fmr1* KO mice have increased AGS susceptibility.

In this study we assessed seizure burden and performed quantitative analysis of the baseline and auditory stimulus-induced EEG activity in *Fmr1* cKO mice. We determined that *Fmr1* cKO mice develop infrequent, brief, spontaneous electrographic seizures at baseline, while WTLM do not exhibit electrographic seizures. This suggests hyperexcitability of neurons under basal conditions in *Fmr1* cKO mice, which is consistent with a recent study that found that the mRNA of the potassium channel Kir4.1 is a target of FMRP and that decreased expression of this channel contributes to neuronal hyperexcitability when *Fmr1* is selectively knocked out in hippocampal astrocytes (Bataveljic et al., 2024). By extension, our results also suggest that lack of FMRP in astrocytes contributes to the increased propensity of seizures in FXS patients. However, neither electrographic nor behavioral seizures occurred in 6-7 week old *Fmr1* cKO mice in response to the auditory stimuli. The absence of siren-induced seizure behavior in adolescent mice and lack of spontaneous seizures in adult mice are not surprising, given that AGS susceptibility rapidly declines in *Fmr1* KO mice older than three weeks (Musumeci et al., 2000; Yan et al., 2004; Yan et al., 2005) and seizures typically resolve in FXS patients before they reach adulthood (Berry-Kravis et al., 2021).

Whisker stimulation provides a translational method for studying response to tactile stimuli. Mechanoreceptors in the whisker follicles of rodents and in the skin of humans can transmit similar types of tactile information to the same brain regions, indicating that whisker stimulation studies can be useful for understanding tactile processing in humans (Diamond, 2010; Juczewski et al., 2016). We did not observe a significant difference between *Fmr1* cKO and WTLM mice in a whisker stimulation paradigm that has been previously used to demonstrate tactile hypersensitivity in *Fmr1* KO mice (He et al., 2017). Although our experiments did not reveal a contribution of loss of astrocytic FMRP to tactile hypersensitivity, it is possible a more intense paradigm, such as the whisker nuisance test in which mice receive bilateral whisker stimulation for a longer period of time (Chelini et al., 2019), would expose genotype differences not detected here.

When compared to previously published studies, our results appear to have discrepancies in two areas. In contrast to the behavior of *Fmr1* KO mice that has been observed in open field assays, in which *Fmr1* KO mice move significantly more than WT controls (Kazdoba et al., 2014; Mineur et al., 2002), we observed significantly <u>less</u> movement in the KO group both during baseline and sham stimulation trials. The main difference between open field tests and our whisker stimulation experiment is that our mice underwent extensive habituation prior to the testing day, making the MHC a familiar environment, whereas in the open field test, mice are placed in a novel environment. Less movement by *Fmr1* KO mice in a familiar environment is consistent with what we have observed during home cage monitoring (Bonasera et al., 2017). In addition, unlike previously reported (He et al., 2017), we found that *Fmr1* KO mice moved for a greater amount of time during whisker stimulation than during sham stimulation, while WTLM do not exhibit this phenotype. This difference could be attributed to the surface on which the mice were running. We used a flat air-lifted cage while the previous study used a polystyrene ball, and it would be expected that mice would exhibit different movement patterns on a flat surface compared to curved surface. One thing to note is that WTLM from *Fmr1* cKO litters tend to behave differently than WTLM from KO litters. This could be attributed to strain differences or the administration of tamoxifen to WTLM from cKO litters but not from KO litters.

Another indication of hypersensitive responses to sensory stimuli is lack of habituation to repeated stimulation. In humans, this can manifest behaviorally as inability to tune out a recurring background sound, such as an air conditioner running, or an enhanced awareness of clothing contacting the skin. Neural correlates of habituation have also been described. FXS patients do not exhibit the typical decrease in amplitude of event related potentials corresponding to repeated auditory stimuli (Ethridge et al., 2016). A similar phenomenon was observed in *Fmr1* KO mice (Lovelace et al., 2016), which also show reduced habituation to repeated whisker stimulation and failure to exhibit adaptation, defined as a reduction in neuronal firing at later stimulations (He et al., 2017). However, we did not observe any behavioral indications of adaptation deficiency in *Fmr1* KO mice, suggesting that detection of adaptation in mice may require more sensitive approaches, such as neuronal imaging or recording.

Our partial conditional knockdown could be considered a disadvantage of the model we used for this study, but in fact, it makes the model relatively translatable, as FXS patients commonly have low residual levels of FMRP (Jacquemont et al., 2018; Loesch et al., 2004; Pieretti et al., 1991). But in order to identify potential therapeutic targets, it will be crucial to obtain a better understanding of the molecular mechanisms underlying these processes. Astrocytes integrate and process information and subsequently modulate neuronal activity through transient elevations in intracellular Ca^2+^ concentrations. This occurs in response to various stimuli, including sensory stimulation (Lines et al., 2020). Ca^2+^ activity in response to activation of glutamatergic or purinergic receptors is altered in astrocytes from *Fmr1* KO mice (Higashimori et al., 2013; Reynolds et al., 2021), as well as in astrocytes differentiated from induced pluripotent stem cells from FXS patients (Ren et al., 2023), suggesting that altered astrocytic Ca^2+^ signaling could contribute to the phenotypes we observed here. Another potential mechanism by which astrocytes lacking FMRP might contribute to sensory hypersensitivity is through dysregulation of inhibitory neuronal signaling. Tactile defensiveness behaviors in response to whisker stimulation can be improved by modulation of inhibitory neuronal signaling either during or after the critical period in *Fmr1* KO mice (Kourdougli et al., 2024; Kourdougli et al., 2023). Our observation that *Fmr1* cKO mice exhibit auditory hypersensitivity suggests that there may be impaired astrocytic regulation of inhibitory neuronal signaling in the absence of FMRP. This could be a result of altered expression or secretion of inhibitory synapse-regulating factors such as neurocan, a proteoglycan secreted by astrocytes that promotes inhibitory synapse formation and is also a target of FMRP (Darnell et al., 2011; Irala et al., 2024). Astrocytes may also contribute to the impaired formation of perineuronal nets around parvalbumin interneurons that has been observed in the auditory cortex of *Fmr1* KO mice (Wen et al., 2018). Aggrecan, a key component of perineuronal nets that is expressed by both neurons and astrocytes, is also a target of FMRP and has reduced translation in the cortex of young *Fmr1* KO mice (Darnell et al., 2011; Van’t Spijker & Richter, 2024).

Here we demonstrated that astrocytic FMRP is critical for normal auditory processing, but may not be involved in tactile processing. However, this could be due to the nature of the stimuli that we used. The auditory stimulation we tested here was a single intense, bilateral, prolonged stimulus, while for tactile stimulation we used a series of short, unilateral, and relatively mild stimuli. Given that astrocytic responses to sensory stimuli, indicated by changes in intracellular Ca^2+^ concentrations, vary depending on the duration, frequency, or intensity of sensory stimuli (Lines et al., 2020; Wang et al., 2006), it could be expected that astrocytes would have differential responses to our different stimuli, and in turn have varying levels of contribution to behavioral responses.

