## Supplementary figures 1-4 for "Astrocytic contribution to auditory hypersensitivity in a mouse model of fragile X syndrome"

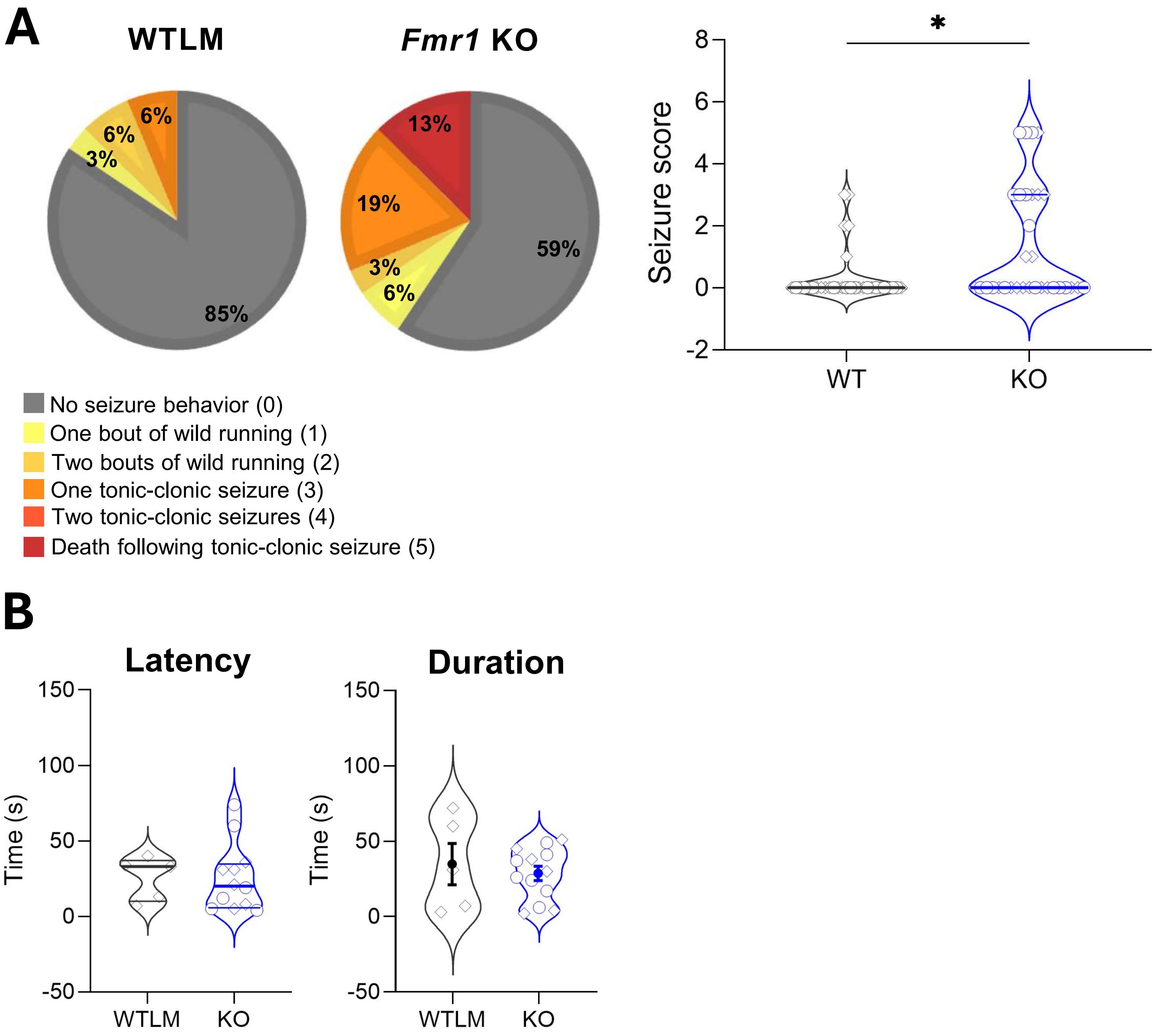


**Supplementary figure 1: *Fmr1* KO mice are susceptible to audiogenic seizures. A)** Proportion of *Fmr1* KO and WTLM mice exhibiting seizure behavior and corresponding seizure score (n=32 WT and 32 KO). **B)** Latency to onset of seizure behavior and total duration of all seizure behavior (n=5 WT and 13 KO). In **A** and **B**, diamonds represent mice that received tamoxifen and circles represent mice that did not. A Chi-square test was used to analyze proportion of mice exhibiting seizures (**A**); Mann-Whitney tests were used to analyze seizure score (**A**) and latency (**B**); unpaired t-test was used to analyze seizure duration (**B**). In **A** and **B** (latency), horizontal lines represent median and upper and lower quartiles. In **B** (duration), filled circles represent mean and bars represent ± SEM. *p < .05

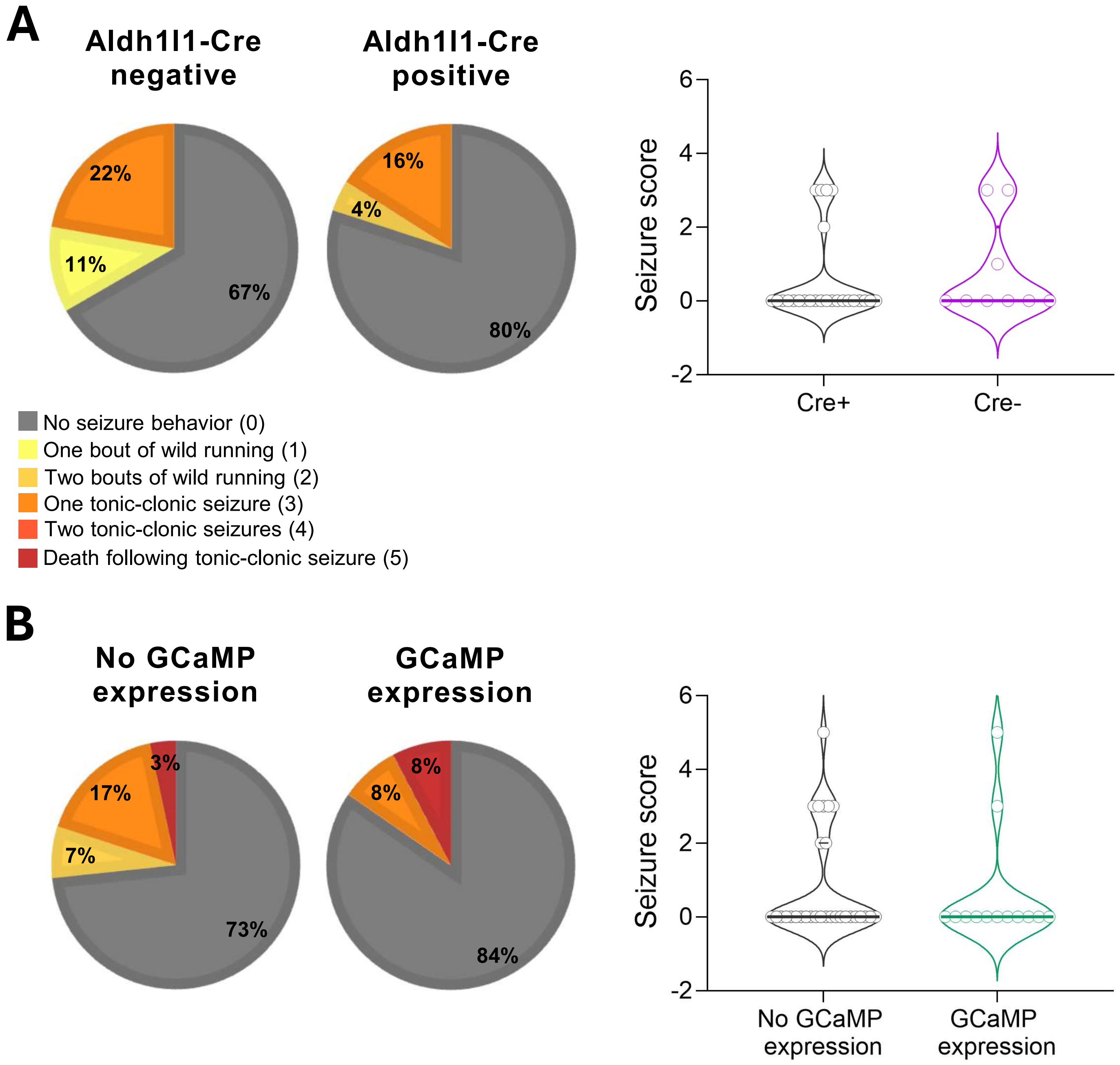


**Supplementary figure 2: Aldh1l1-cre and GCaMP6f expression do not impact AGS susceptibility. A)** Proportion of Aldh1l1-Cre negative and Aldh1l1-Cre positive mice exhibiting seizure behavior and corresponding seizure score (n=9 Cre- and 25 Cre+). **B)** Proportion of mice without or with GCaMP6f expression in astrocytes that exhibit seizure behavior and corresponding seizure score (n=30 without GCaMP expression and 13 with GCaMP expression). Chi-square tests were used to analyze proportion of mice exhibiting seizures; Mann-Whitney tests were used to analyze seizure score. Horizontal lines represent median and upper and lower quartiles.

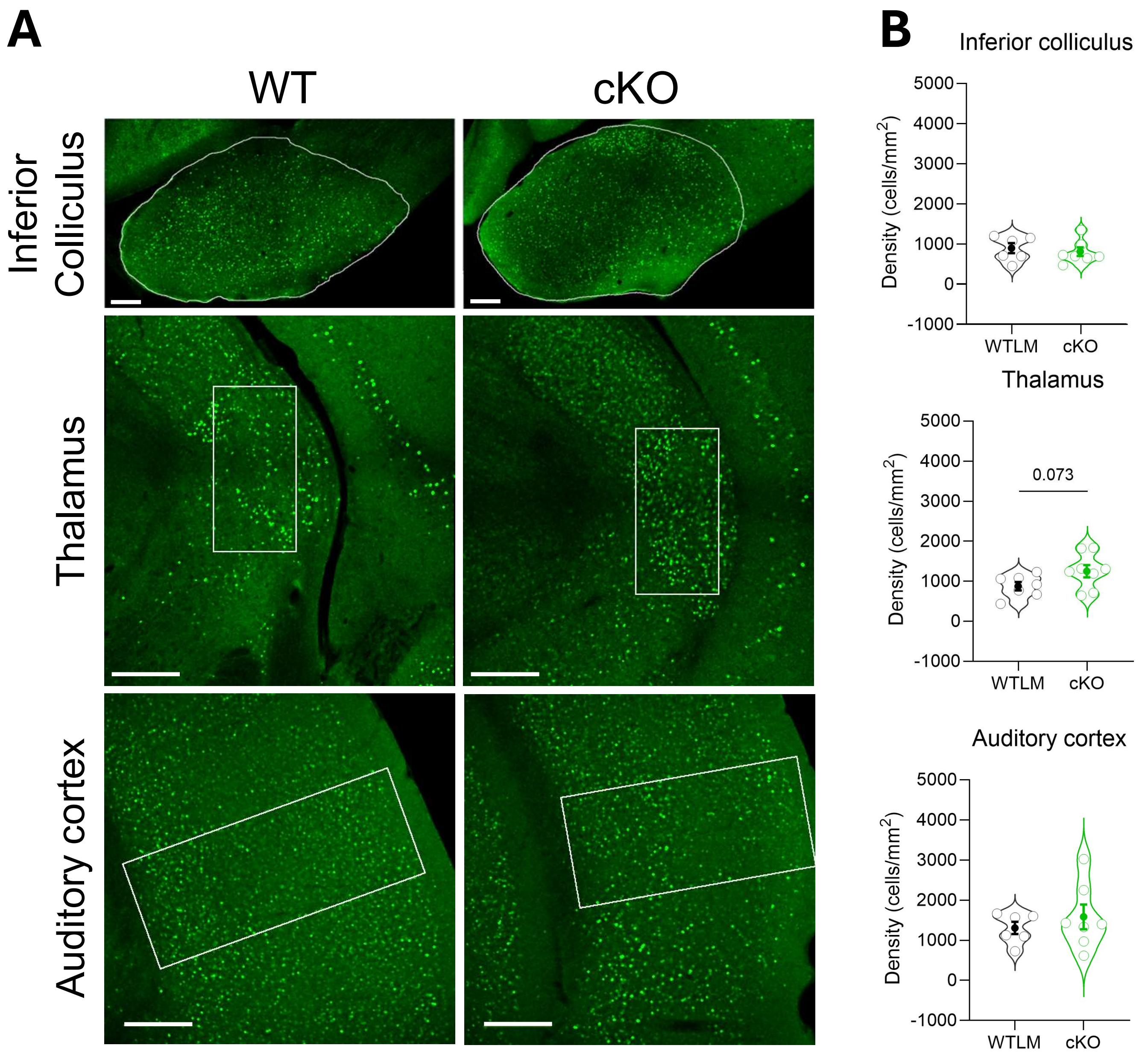


**Supplementary figure 3: Neuronal activation following sound exposure.**
**A)** Representative confocal images of c-Fos immunolabeled sections from the auditory processing pathway. Region analyzed is outlined in white. Scale bar is 200 µm. **B)** Density of c-Fos+ cells in the respective brain regions (n = 6-8 mice, average of 2-3 images/mouse). Unpaired t-tests were used to analyze density of c-Fos+ cells. Filled circles represents mean and bars represent ± SEM.

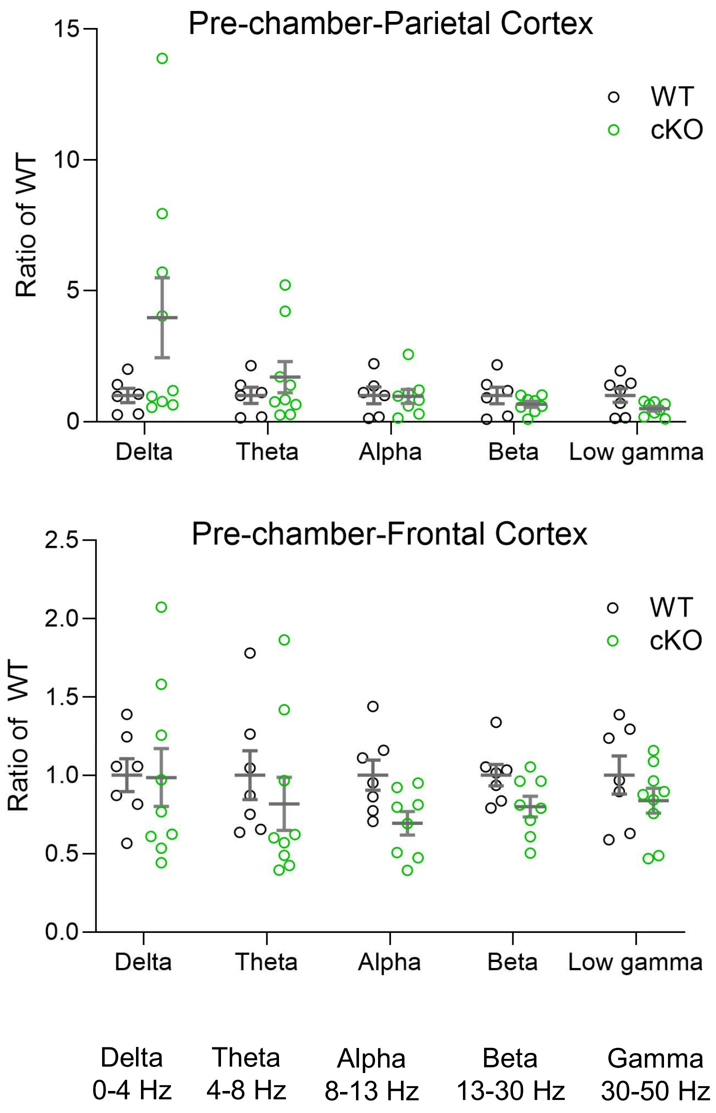


**Supplementary figure 4: Neural oscillations are not altered in *Fmr1* cKO mice.** Average power of neural oscillations in the parietal cortex and frontal cortex of *Fmr1* cKO mice and WTLM. Average power values are displayed as a ratio of WT for each frequency range. Two-way ANOVA tests were used to analyze differences between genotypes and frequencies.
